# Viral vectors with cluster of differentiation gene promoters to target specific cell types in the brain

**DOI:** 10.1101/2024.10.21.619372

**Authors:** Naofumi Uesaka, Mariko Sekiguchi, Atsuya Takeuchi, Ryo Masumura

**Author notes:** lead contact, Naofumi Uesaka, Department of Cognitive Neurobiology, Graduate School of Medical and Dental Sciences, Institute of Science, Tokyo, 1-5-45 Yushima, Bunkyo-ku, Tokyo 113-8510, Japan.

## Abstract

Understanding brain function and developing targeted therapies for neurological disorders require precise access to specific cell types, but current methods are limited. Here, we report the development of cell-type specific adeno-associated virus (AAV) vectors utilizing promoters of cluster of differentiation genes (CD promoters) for targeted gene delivery in the brain. We newly identified CD promoters that induce gene expression selectively in specific cell types in the mouse brain. These AAVs enable in vivo calcium imaging and chemogenetic applications in specific cerebellar cells, revealing distinct and shared roles of two types of cerebellar interneurons. Notably, chemogenetic modulation of cerebellar molecular layer interneurons using a CD promoter rescued social behavior and motor deficits in a mouse model of autism spectrum disorder. Our findings demonstrate the utility of these AAVs in elucidating the functions of individual cell types in brain function and in developing novel, cell-type specific therapies for brain diseases.

## Introduction

One of the fundamental quests in the field of neuroscience is to define the role of each cell in brain function and neural circuits. This knowledge is key to unlocking how each cell generates complex brain functions and how dysfunction of each cell results in various brain diseases. Achieving this goal necessitates defining each cell type and generating genetic tools that can label and manipulate in a highly cell-selective manner. Recent technological advances such as single-cell RNA sequencing have facilitated cataloguing gene expression profiles among different populations of cells in the nervous system and have defined each cell type based on genome-wide gene expression, thereby enabling identification of diverse cellular subtypes in the brain.

Even with these technological advances, there are considerable limitations. Most notably, the current set of tools does not permit individual access to most of cell types present in the brain. Recent innovations in viral technology offer promising avenues for overcoming the obstacle. Viral vectors such as adeno-associated virus (AAV) vectors are versatile and can deliver any genes into neurons and glial cells across different species. Additionally, AAVs are emerging as potent clinical tools for treating a wide range of diseases, as AAVs can specifically target and genetically manipulate specific mutant cells, thereby enabling the treatment of a wide range of diseases. Thus, the construction of AAV-based transgenic tools is useful in many fields. Current researches have shown that multiple elements like serotypes, promoters, enhancers, and microRNA targeting sequences in AAVs can affect the efficiency of infection and gene expression in specific cell types. Although cell type-specific AAVs with these elements have been partially developed ^1–6^, AAVs that can target specific cell types is limited.

To overcome this limitation, we developed cell-type-specific AAVs using promoters of cluster of differentiation (CD) genes. CD numbers, assigned to specific cell surface antigens, help identify fine cell type differences, and various CD genes are expressed in the brain, offering unique targeting opportunities ^7–10^. We systematically screened AAVs with each CD gene promoter (CD promoter) which induced the expression specifically in each type of cell in the cerebellum and found some CD promoters enabled specific expression in each type of cell in the cerebellum. We revealed three CD promoters which expressed a gene of interest specifically in excitatory neurons and parvalbumin positive inhibitory neurons in the mouse cortex, indicating that our approach was not only successful in the cerebellum but also in the cortex. Using the AAVs with CD promoters, we revealed that two types of cerebellar interneurons played distinct roles in social behavior, locomotor activity, and motor function, and the shared role in anxiety behaviors. Additionally, the selective activation of cerebellar molecular layer interneurons by AAVs with the specific CD promoter rescued social deficits in a mouse model of autism spectrum disorder. The development of cell-type-specific AAV vectors utilizing CD promoters enables targeted manipulation of neurons that were previously difficult to access, such as cerebellar granule cell layer interneurons, providing new insights into their roles in brain function and offering novel therapeutic strategies for neurological disorders.

## Results

### Screening of CD promoters in the mouse cerebellum

To identify CD promoters which enable gene expression in specific type of cells, we focused on the mouse cerebellum (Figure. 1A) because AAVs that can target specific cell types are limited compared with other nervous system. We detected 31 CD promoters that are expressed in the cerebellum from our previous microarray data ^11^. We amplified approximately −700 bp to −2200 bp from the transcriptional start site of each CD promoter using the mouse and the human genomes by PCR, and then cloned these promoter sequences into an AAV backbone with mcherry or EGFP reporter (Figure. 1B, Table 1). Viral particles were packaged with the AAV9 capsid and injected into the cerebellum of adult mice (Figure. 1B). After 12–14 days, we fixed the brains and performed immunohistochemistry.

**Figure. 1:**
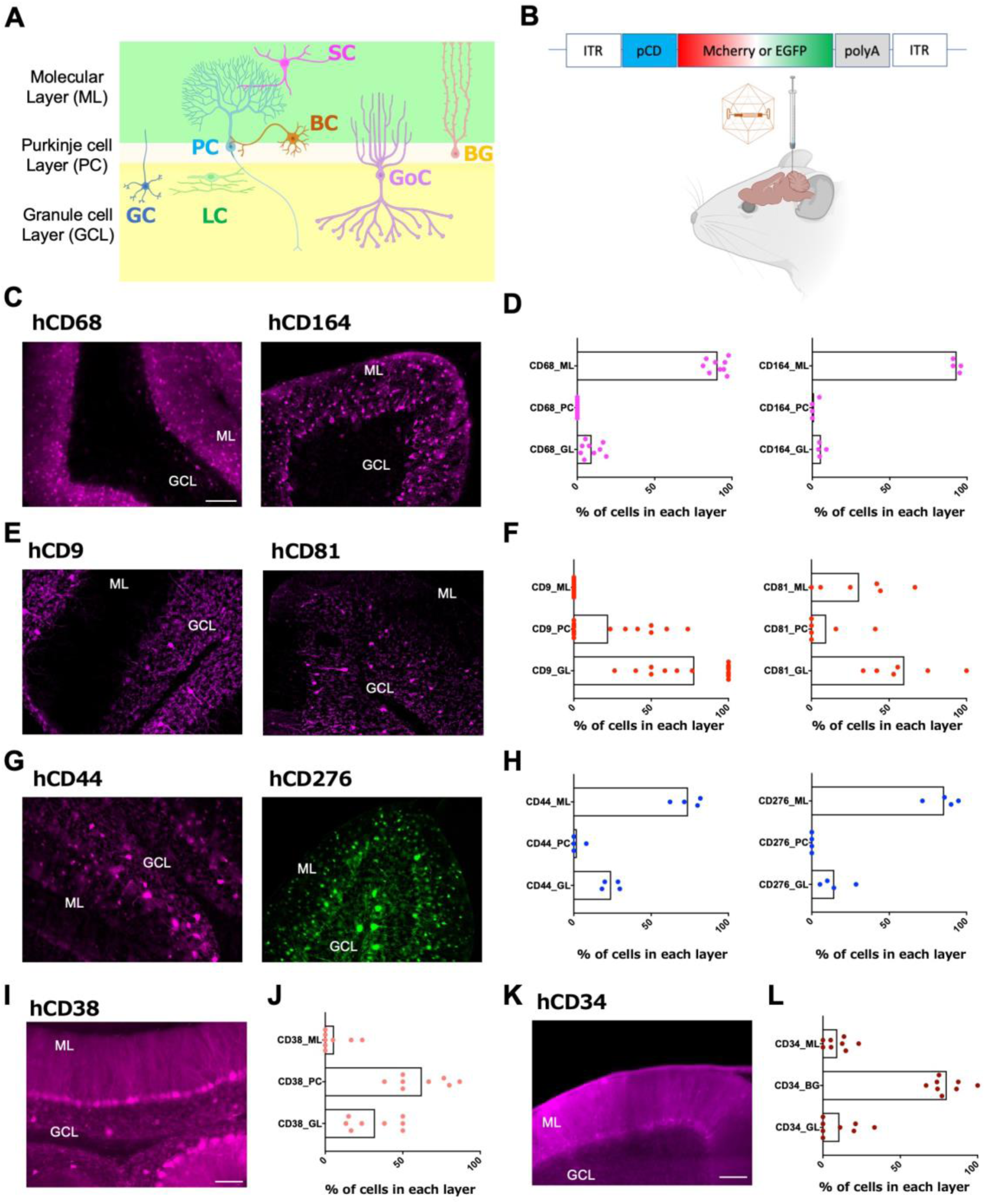
CD promoters drive gene expression in specific type of cells in the mouse cerebellum. **A**, layers and cell types in the mouse cerebellum. **B**, Schematic illustrating the expression cassettes of the AAV vectors with CD promoters and AAV injection into the mouse cerebellum. **C,** mcherry fluorescent images of the cerebellum infected by AAVs with one of hCD68 and hCD164 promoters. **D**, graphs showing the percentage of mcherry^+^ cells in each cerebellar layer for hCD68 and hCD164 promoters. n = 2 mice for each. **E-L,** similar fluorescent images and graphs to c,d for hCD9 and hCD81 promoters (e, f), for hCD44 and hCD276 promoters (g,h), hCD38 promoter (i, j), hCD34 promoter (k, l). For hCD276, the EGFP image (g) and the percentage of EGFP^+^ cells in each cerebellar layer (h) are shown. Scale bar, 100µm. GC, granule cell. PC, Purkinje cell. LC, Lugaro cell. BC, basket cell. SC, stellate cell. GoC, Golgi cell. BG, Bergman glia. ML, molecular layer. PC, Purkinje cell. GL or GCL, granule cell layer. BG, Bergman cell.

**Table 1.**
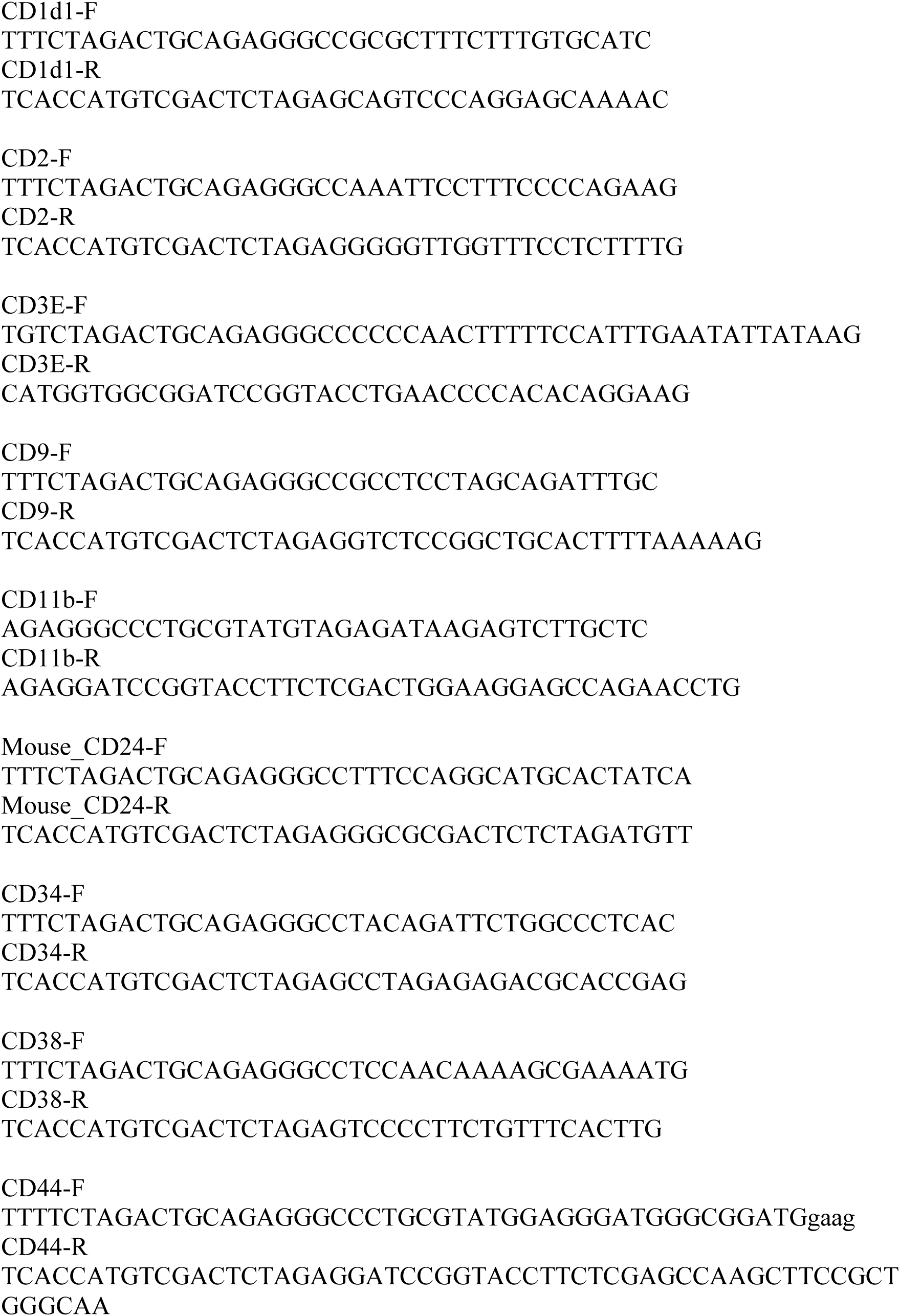
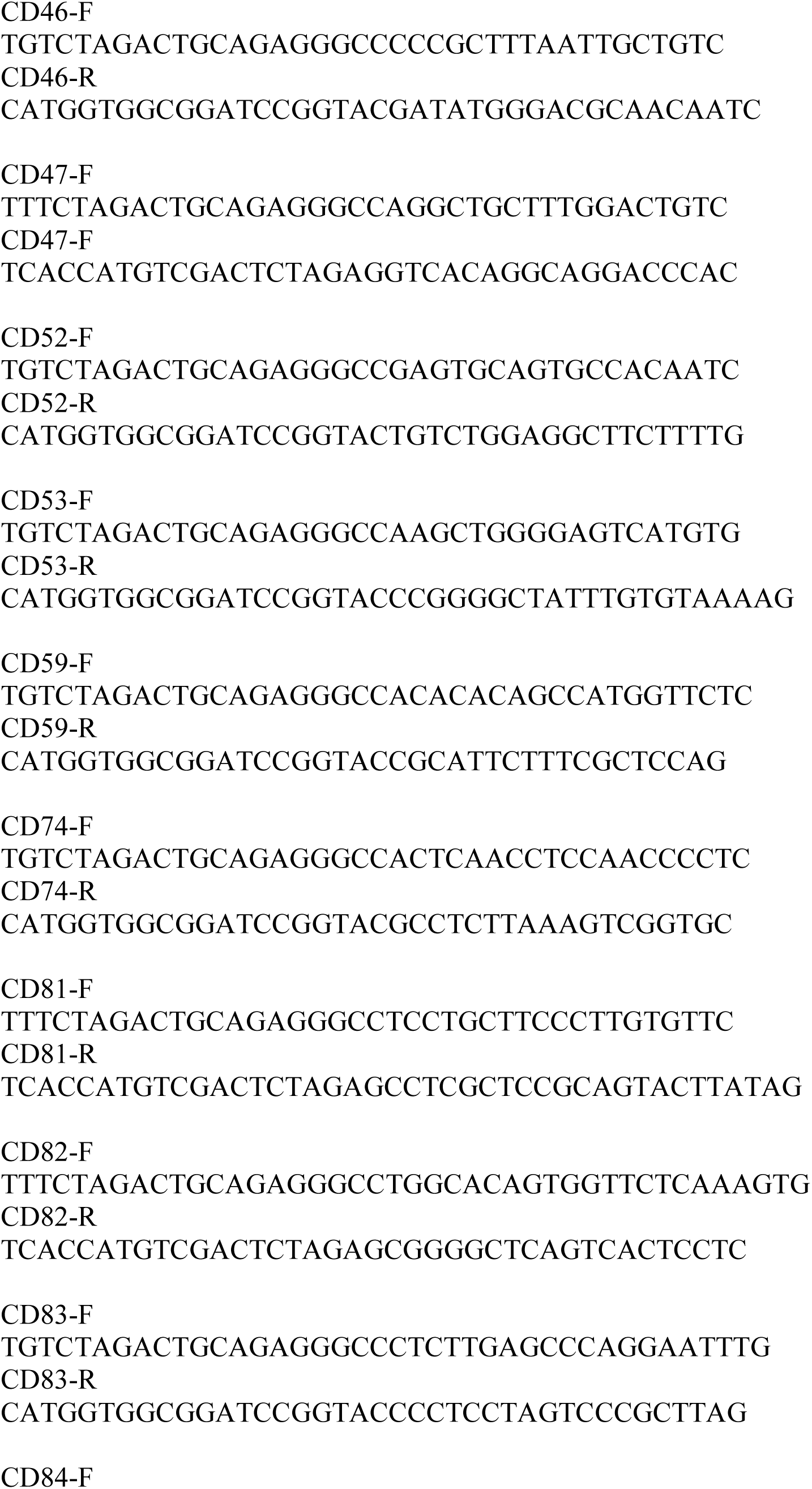
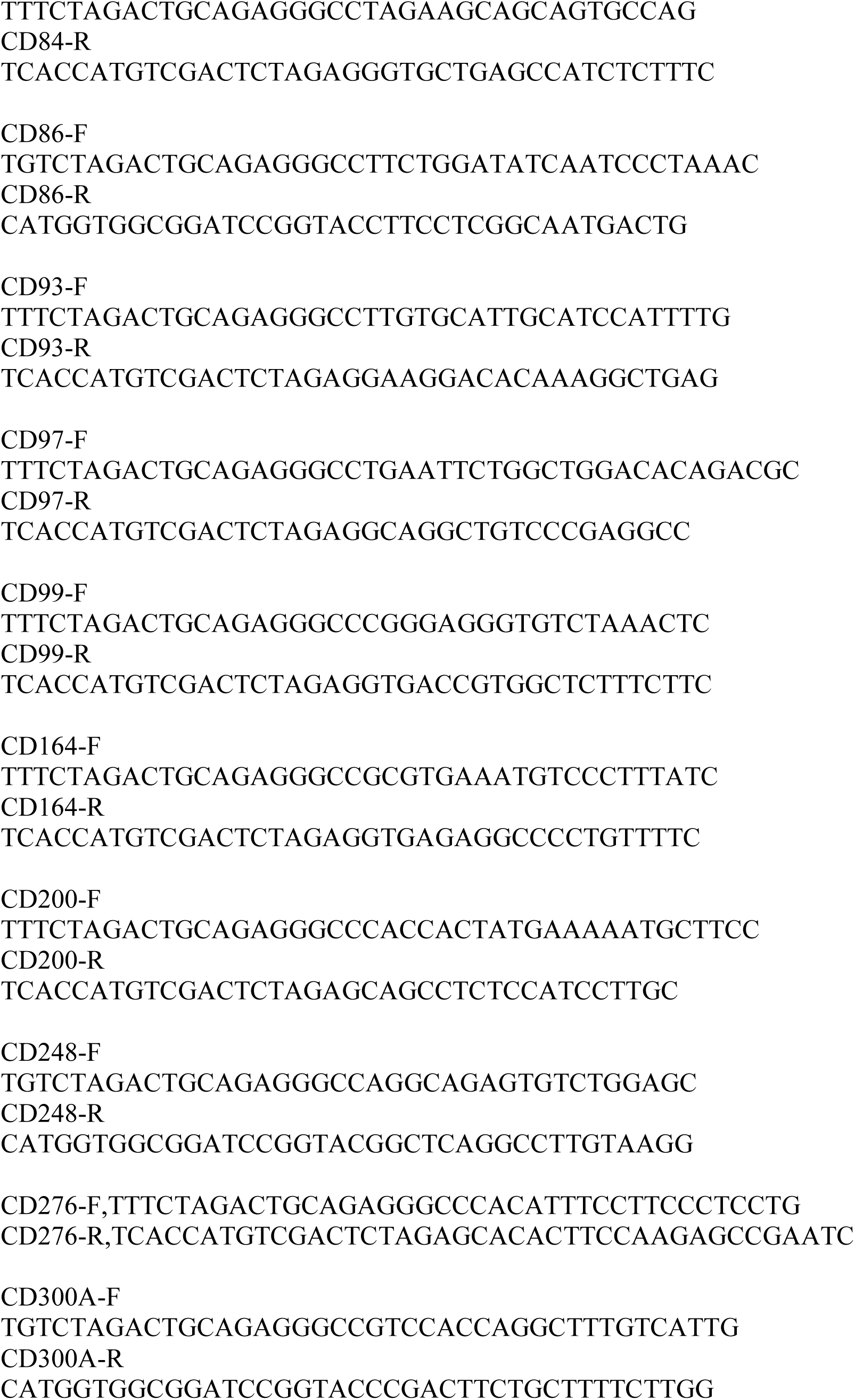
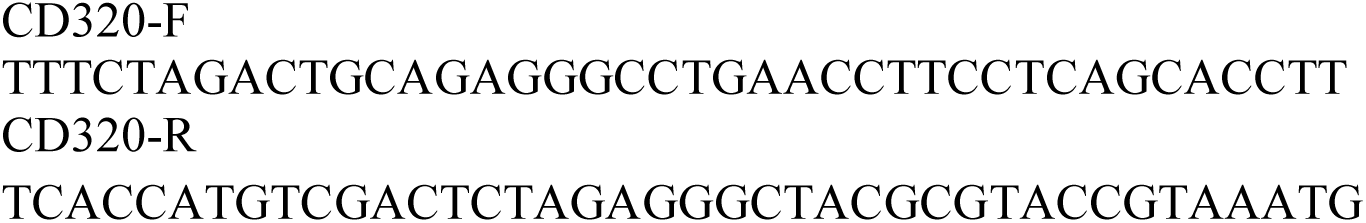
PCR primer sequences of CD promoters (5’-3’)

All AAVs with CD promoters drove expression of the reporter gene in the cerebellum. The grey matter of the cerebellar cortex is divided into three layers: the outer molecular layer (ML), the middle Purkinje cell layer (PCL), and the inner granule cell layer (GL) (Figure. 1A), each containing different cell types. To determine which cerebellar layers were targeted by each CD promoter, we analyzed the location of reporter-labeled cells. We found that AAVs with human CD52 (hCD52), hCD68, hCD74, hCD83, and hCD164 promoters mainly drove gene expression in the ML, with sparse activity in the GL (Figures. 1C-D and S1A-B). In contrast, the vast majority of cells labeled by the AAV with the hCD9 promoter were in the GL, where Golgi cells appeared to be labeled, with a small fraction of labeled cells being Purkinje cells (Figure. 1E-F). Similarly, with the hCD81 promoter, Golgi cells were predominantly labeled, but some labeled cells were present in the ML and PCL. AAVs with either hCD44 or hCD276 promoters labeled cells in both the molecular layer and the granule cell layer (Figure. 1G-H). The hCD300 promoter drove EGFP expression in the ML and in the GL where Lugaro cells seemed to be labeled (Figure. S1C-E). Most of the cells labeled by the AAV with the hCD38 promoter were Purkinje cells, with additional labeled cells present in the GL where Golgi cells appeared to be labeled (Figure 1I-J). The hCD34 promoter was quite selective for cells expressing S100β, a marker for Bergmann glia, specialized astrocytes in the cerebellum (Figures. 1K-L and S1F).

To address whether the observed layer-specific expression was due to promoter specificity rather than differences in cell distribution, viral spread, or infection efficiency, we compared the expression patterns of the CD promoter-driven AAVs with that of an AAV using the pan-neuronal synapsin promoter driving mCherry expression. The synapsin promoter induced widespread expression across all cerebellar layers with relatively uniform distribution of labeling neurons in the ML, PCL, and GL (Figure. S2). This distribution suggests that our injection method effectively delivers the virus throughout the cerebellum and that the number of PV+ neurons in the ML being greater than in the GL does not account for the specific expression patterns observed with the CD promoters. Therefore, these data indicate that specific CD promoters have intrinsic preferences for regulating gene expression in distinct layers and cell types within the mouse cerebellum.

### Cell-type specificity enhanced by combinations of promoters, enhancers, and microRNA targeting sequences

To scrutinize and optimize the cell-type specificity of our CD promoter-based AAV vectors, we focused on the hCD68 and hCD9 promoters. We used the immunoreactivity for parvalbumin (PV) as a marker for molecular layer interneurons and neurogranin (NG) as a marker for Golgi cells. We found that the hCD68 promoter selectively labeled PV^+^ molecular layer interneurons (>94 %) in the ML (Figure. 2A and 2E), indicating minimal labeling of other cells in the ML. However, a small fraction of cells in the GL were also labeled (Figure. 1D and 2D). The hCD9 promoter enabled specific labeling of Golgi cells that existing vectors cannot label (Figure. 2F and 2K), but a small fraction of cells in the PCL were also labeled (Figure. 1F and 2I).

**Figure. 2:**
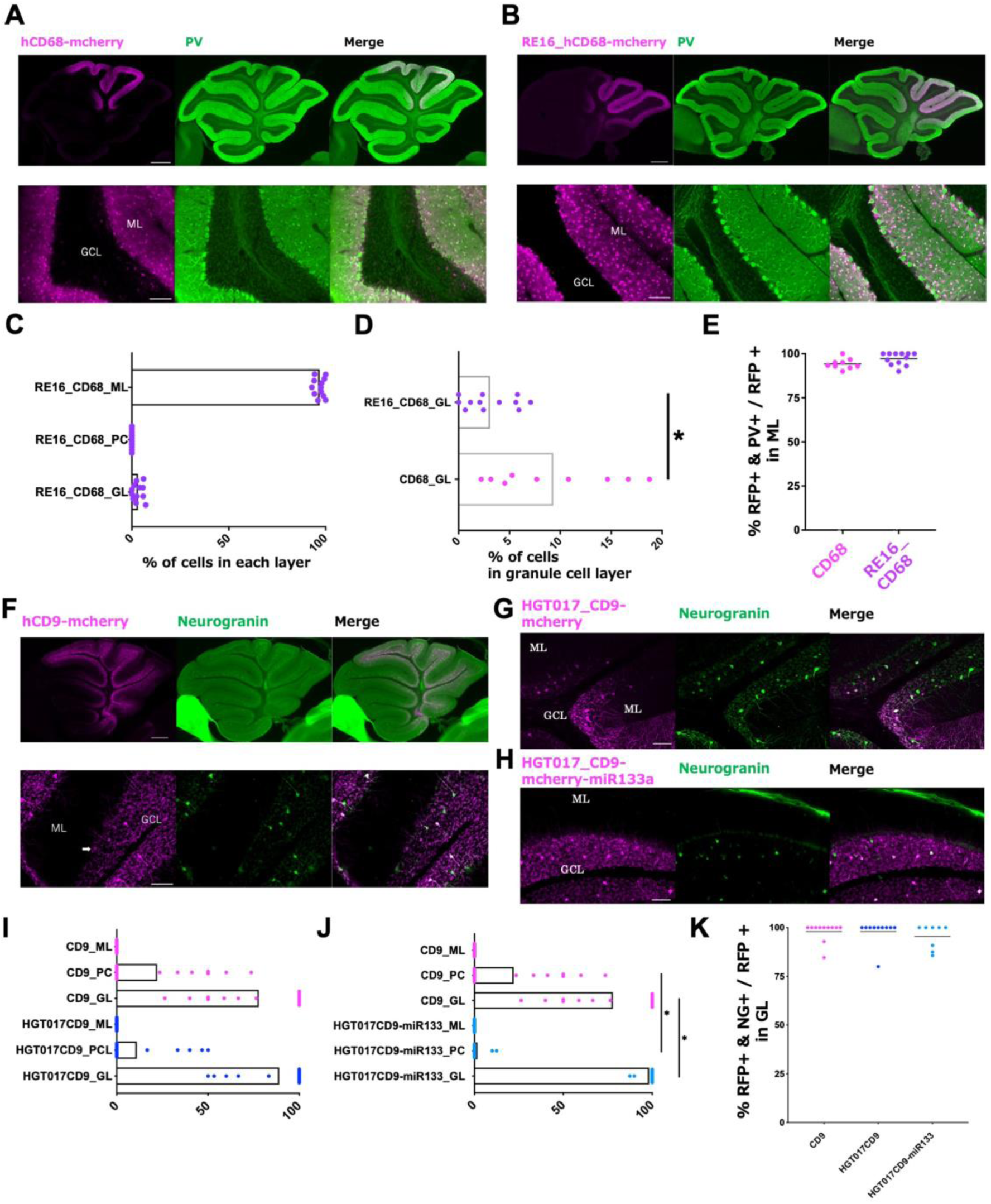
The combination of CD promoters with enhancers enhances the specificity to each type of cell in the mouse cerebellum. **A**, fluorescent images of mcherry (magenta) and PV (green) in the cerebellar slice infected by AAVs with mcherry under the hCD68 promoter. Bottom, the magnified image of the top image. **B**, the images for hCD68 promoter with the mscRE16 enhancer similar to A. **C**, the graph showing the percentage of mcherry^+^ cells in each cerebellar layer for hCD68 with the mscRE16 enhancer. n = 3 mice. **D**, the graph showing the comparison of the percentage of mcherry^+^ cells in the granule cell layer between the hCD68 promoter with and without mscRE16. **E,** the graph showing the specificity of gene expression in PV^+^ molecular layer interneurons by the hCD68 promoter with/without mscRE16 enhancer. The percentages were determined by the ratio of double-labeled cells (PV^+^ and mcherry^+^) to total mcherry^+^ cells. **F-G**, images and graphs for hCD9 promoter similar to a-e. **F-H**, images (mcherry and neurogranin for low and high magnification) for AAVs with mcherry under the hCD9 promoter (F), hCD9 promoter with the HGT017 enhancer (G), and hCD9 promoter with the HGT017 enhancer and miR133a (H). Arrows show mcherry-labeled Purkinje cells. **I,** the graph showing the percentage of mcherry^+^ cells in each cerebellar layer for hCD9 with or without the HGT017 enhancer. n = 3 mice for each. **J,** the graph showing the percentage of mcherry^+^ cells in each cerebellar layer for hCD9 with or without the HGT017 and miR133a. n = 3 mice for each. **K**, the graph showing the specificity of gene expression in neurogranin^+^ interneurons by the hCD9 promoter with/without HGT017 enhancer and miR133a. The percentages were determined by the ratio of double-labeled cells (neurogranin^+^ and mcherry^+^) to total mcherry^+^ cells. Scale bar, 500µm for top in a, b, f, 100µm for the others. ML, molecular layer. PC, Purkinje cell. GL or GCL, granule cell layer. * p<0.05 (Mann-Whitney U test).

To further enhance the specificity of the hCD68 promoter for molecular layer interneurons, we explored the potential of combining the hCD68 promoter with an enhancer. Several previous studies have identified enhancer sequences for cell-type specific gene expression^1, 6, 12^. We selected a range of enhancers and analyzed the expression pattern of a reporter gene regulated by these enhancers in the cerebellum. The mscRE16 enhancer^1^ drove the expression of the reporter gene mainly in molecular layer interneurons and slightly in granule cell layer (Figure. S3A-D). To consolidate these findings, we created a new AAV construct that combined both hCD68 promoter and mscRE16 enhancer (AAV-RE16_CD68-mcherry). To enhance gene expression, we used AAV-PHP.eB^13^. When we used the similar concentration for both AAV-CD68-mcherry and AAV-RE16_CD68-mcherry, the AAV-RE16_CD68-mcherry resulted in similar infection efficacy (Figure. S3E), but fewer labeled cells in the GL compared to AAVs using just the hCD68 promoter and mscRE16 enhancer (Figure. 2B-E and S3C).

We further investigated whether systemic delivery of AAV-RE16_CD68-mCherry via retro-orbital injection could specifically target molecular layer interneurons in the cerebellum. We employed retro-orbital injection as a less invasive alternative to direct intracerebellar administration (Figure. S3F). Retro-orbital injection of AAV-RE16_CD68-mCherry resulted in similar specificity for cerebellar molecular layer interneurons as observed with direct intracerebellar injection (Figure. S3G-H). These findings suggest that the cell selectivity of AAV-RE16_CD68-mCherry is not dependent on the injection site and is intrinsic to the vector itself. Therefore, retro-orbital injection can serve as an effective and less invasive alternative for targeting cerebellar molecular layer interneurons using AAV-RE16_CD68-mCherry.

To enhance the specificity of AAVs with hCD9 promoter, we combined the hCD9 promoter with an enhancer. We used AAV.CAP-B10 as an AAV capsid ^14^ and created a new AAV construct with both the hCD9 promoter and the HGT_017 enhancer^12^. When we used similar concentrations for both AAV-hCD9-mcherry and AAV-HGT017CD9-mcherry, pairing the hCD9 promoter with HGT_017 enhancer resulted in similar infection efficacy (Figure. S4G), but significantly greater targeting specificity to cells in the GL compared to using the hCD9 promoter alone (86.0 ± 6.3 % for HGT017_hCD9, 77.9 ± 6.9 % for hCD9, Figure. 2G, 2I). Additionally, the specificity to Neurogranin^+^ Golgi cells was high for hCD9 and hCD9 with HGT_017 (> 97%, Figure. 2K).

We further attempted to reduce the percentage of labeling cells in PL for the AAV with hCD9 promoter. We employed microRNA (miRNA) targeting sequences aimed at downregulating gene expression in Purkinje cells. We searched miRNAs expressed in Purkinje cells that have been previously reported and detected candidate miRNAs (miR-133a, miR-1188, miR-1983, miR-3086-5p)^15^. We engineered AAV vectors that incorporated the hCD9 promoter, the HGT017 enhancer, and four copies of each miRNA targeting sequence (Table 2). The AAV-HGT017_hCD9-mcherry vector with the miR-133a targeting sequence (miR-133a) induced similar infection efficacy (Figure. S4G), but effectively reduced the percentage of Purkinje cells which expressed mcherry and showed high levels of specificity to cells in granule cell layer (Figure. 2H, 2J). The AAV-HGT017_hCD9-mcherry with other miRNA targeting sequences did not alter percentages of Purkinje cells and cells in the GL (Figure. S4A-F). Immunohistochemical analysis using neurogranin antibody showed that 95.5 % of cells in the GL labeled by mcherry for the AAV-HGT017_hCD9-mcherry with miR-133a was neurogranin^+^ Golgi cells (Figure. 3K). These results suggest that the miR-133a targeting sequence enhances the specificity to Golgi cells by reducing the gene expression in Purkinje cells. Retro-orbital injection of AAV-HGT017_hCD9-mcherry with miR-133a resulted in similar specificity for cerebellar granule cell layer interneurons as observed with direct intracerebellar injection (Figure. S4H-I). These findings suggest that the cell selectivity of AAV-HGT017_hCD9-mcherry with miR-133a is not dependent on the injection site.

**Fig. 3:**
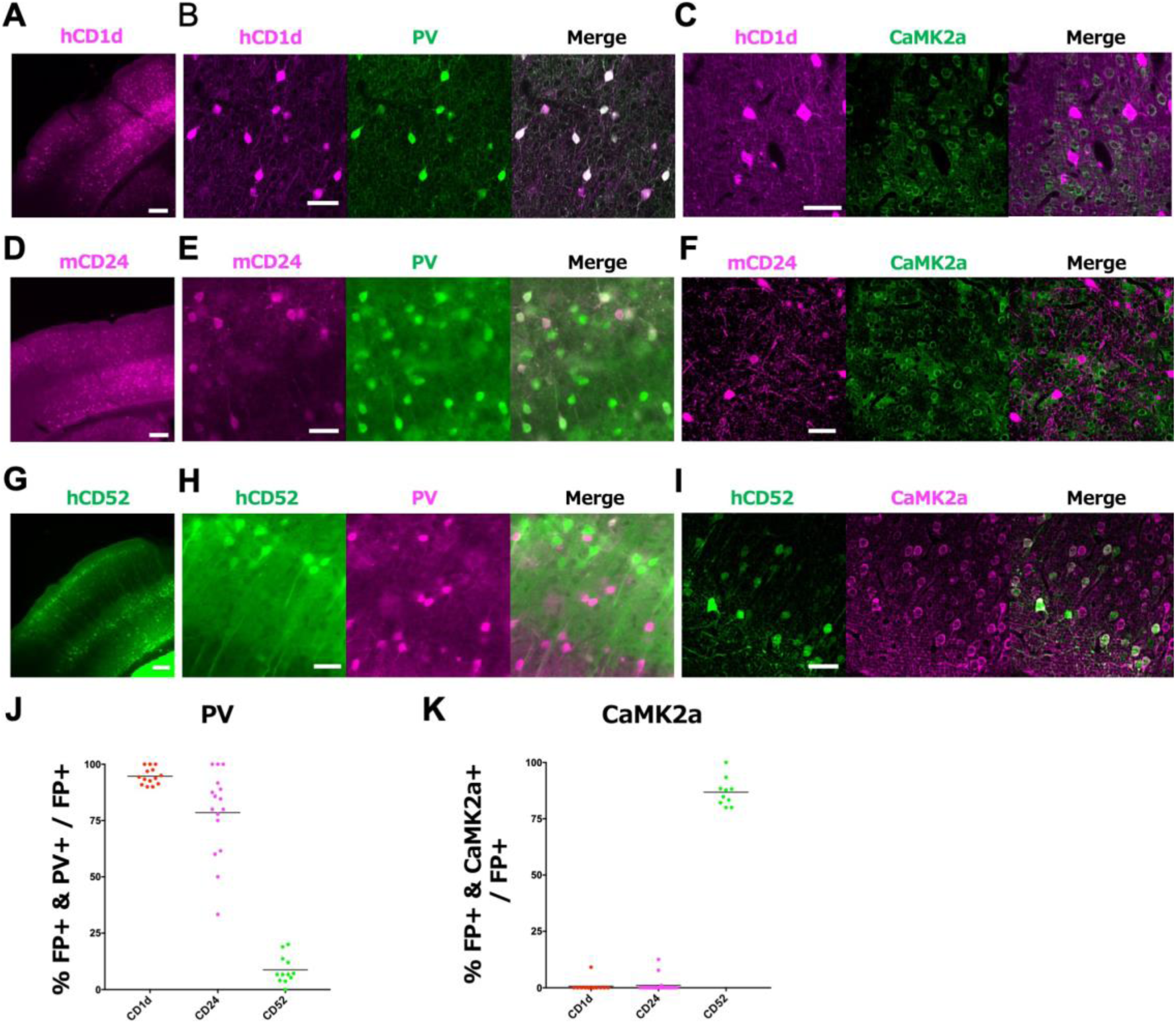
CD promoters drive gene expression in specific type of cells in the mouse cortex. **A**, a mcherry fluorescent image of the mouse cortex infected by AAVs with hCD1d promoter. **B,C**, high magnification images showing mcherry (magenta) and PV (green) for b, and mcherry (magenta) and CaMK2a (green) for c in the cortex. **D-I,** similar fluorescent images to a-c for mCD24 (d-f) and hCD52 (g-i) promoters. For hCD52, EGFP images are shown. **J**, the graph showing the specificity of gene expression in PV^+^ interneurons in the cortex by hCD1d, mCD24, and hCD52 promoters. The percentages were determined by the ratio of double-labeled cells (PV^+^ and mcherry^+^ or EGFP^+^) to total mcherry^+^ cells or EGFP^+^ cells. n = 3 mice for each. **K**, the graph showing the specificity of gene expression in CaMK2a^+^ neurons in the cortex by hCD1d, mCD24, and hCD52 promoters. The percentages were determined by the ratio of double-labeled cells (CaMK2a^+^ and mcherry^+^ or EGFP^+^) to total mcherry^+^ cells or EGFP^+^ cells . n = 3 mice for each. Scale bar, 200µm for a, d, g, 50µm for the others.

**Table 2.**
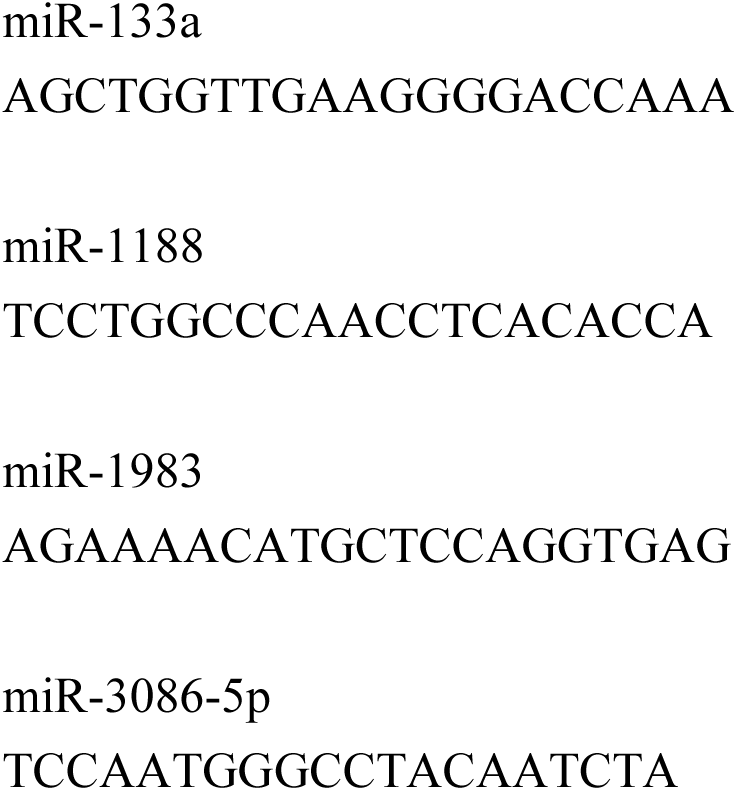
microRNA targeting sequences.

### Identification of cell-type specific CD promoters in the mouse cortex

We next evaluated the scalability of CD promoters by archiving cell-type specific expression in the mouse cortex. Although AAV vectors utilizing promoters and enhancers have been employed to target inhibitory and excitatory neurons ^1, 3, 4, 6, 16^, expanding the repertoire of available promoters enables researchers to select the most appropriate DNA element to achieve desired expression levels, temporal control, or to overcome limitations posed by the cellular environment in different disease models. Using AAVs with CD promoters as those screened in the cerebellum, we tried to identify CD promoters which target inhibitory neurons and excitatory neurons in the mouse sensory cortex. We injected AAVs with CD promoters directly into the mouse cortex and evaluated cell-type specificity through immunohistochemical assays utilizing PV (inhibitory marker) and CaMK2a (excitatory marker) antibodies.

By screening 31 CD promoters, we identified three types of CD promoters (hCD1, mCD24, and hCD52) which demonstrated cell-type specificity in the mouse cortex. Injection of AAV with hCD1d promoter resulted in selective labeling of PV^+^ cortical interneurons with a specificity exceeding 94% (Figure. 3A, 3B, 3J). Moreover, the AAV drove minimal gene expression in CaMK2a^+^ cortical excitatory neurons (Figure 3C, 3K), indicating that the hCD1d promoter works specifically in cortical PV^+^ inhibitory neurons. The AAV with mouse CD24 (mCD24) promoter showed 78.5 % specificity toward PV^+^ neurons of the mouse cortex, with negligible expression in CaMK2a^+^ excitatory neurons (<1 %), suggesting that mCD24 is selective for PV^+^ neurons and other types of inhibitory neurons (Figure. 3D-F, 3J-K). Finally, the CD52 promoter drove the EGFP expression with over 86 % specificity in CaMK2a^+^ excitatory neurons of the cortex, but minimal expression in PV^+^ neurons (<9 %) (Figure. 3G-K). These findings suggest that CD promoters can be effectively employed to drive cell-type-specific gene expression in both the cerebellum and the cortex.

### Intravital calcium imaging of cerebellar Golgi cells

We explored the utility of AAV vectors containing CD promoters for studies aimed at detecting neural activity patterns, focusing on cerebellar Golgi cells—an inhibitory interneuron population located deep within the granule cell layer. Performing in vivo calcium imaging of Golgi cells using a single AAV vector has been challenging with existing techniques due to the requirement of the generation of transgenic mice expressing calcium indicators specifically in Golgi cells, a process that is both time-consuming and costly. Using AAV vectors with the HGT017_hCD9 and the miR133a_targeting sequence, we induced the expression of jGCaMP8s in cerebellar Golgi cells. One week after AAV injection, jGCaMP8s signals were selectively observed in Golgi cells expressing neurogranin (Figure. 4A). These jGCaMP8s signals were absent in Car8-positive Purkinje cells and in molecular layers (Figure. 4A), thus confirming the specificity of jGCaMP8s expression for Golgi cells. We performed in vivo calcium imaging one week after AAV injection and found that Golgi cells expressing jGCaMP8s displayed marked fluorescence changes, showing spontaneous Ca^2+^ transients (Figure. 4B, 4C). The ability to monitor the activity of Golgi cells in vivo without relying on transgenic mice provides valuable insights into their functional roles in cerebellar processing, which has been difficult to achieve with previous methods.

**Figure. 4:**
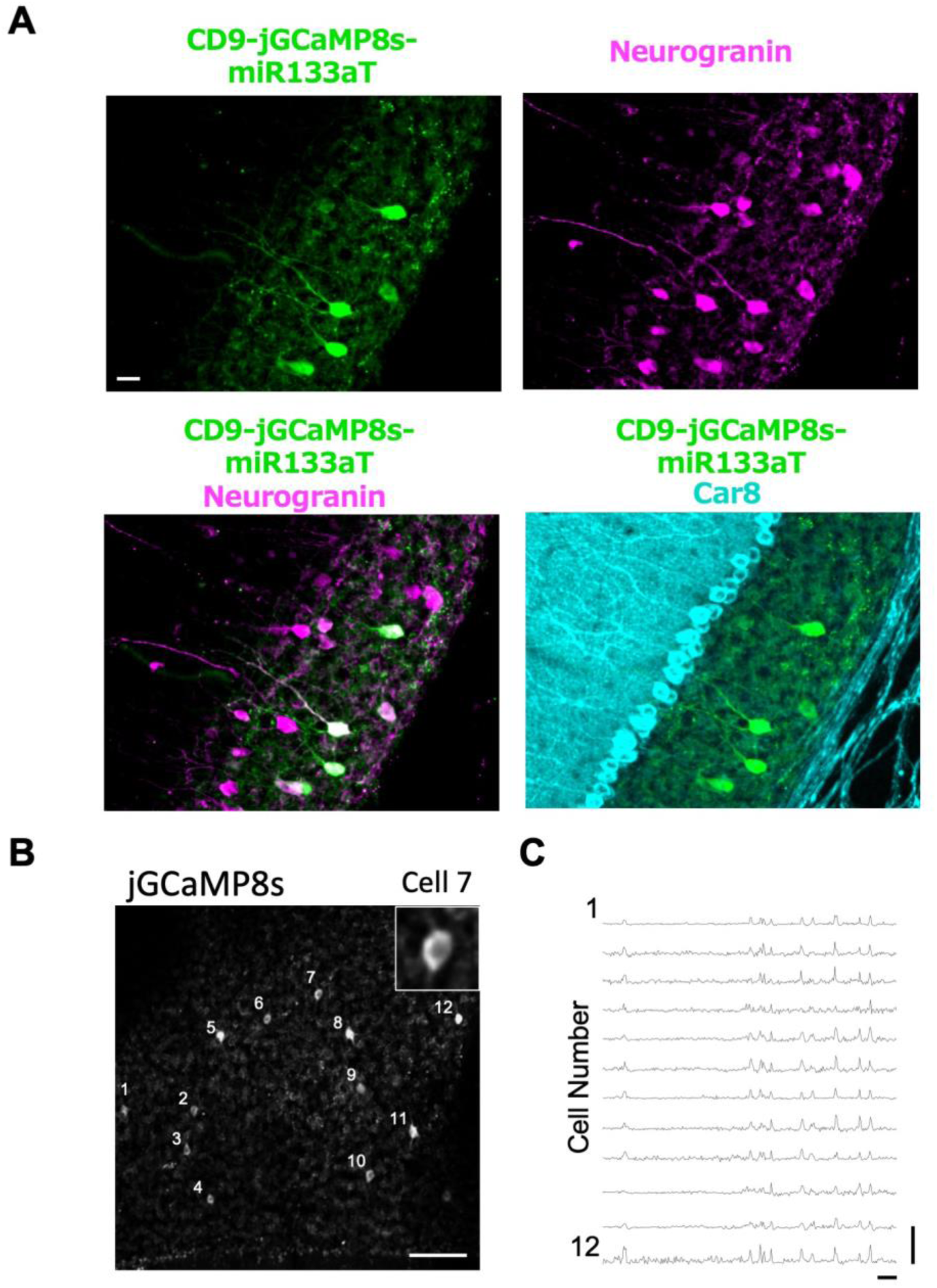
Intravital calcium imaging for neurogranin^+^ Golgi cells in the cerebellum. **A,** fluorescent images of jGCaMP8s (green) and neurogranin (Golgi cells, magenta) or Car8 (Purkinje cells, cyan) in the cerebellar slices infected by AAVs with jGCaMP8s under the hCD9 promoter with the HGT017 enhancer and the miRNA133a targeting sequence. Scale bar, 20 µm. **B**, full-frame scan movies recorded at the granule cell layer at postnatal 9 weeks. Analyzed cells were numbered (in white). Scale bar, 100 µm. Insert, the higher magnification image for cell no. 7. **C**, calcium transients by extracting relative fluorescence changes from each ROI in the panel. Scale bar, 20 s, 10 ⊿F/Fb.

### Modulation of behaviors by manipulation of specific cell types in the cerebellum utilizing CD-promoter-based AAVs

To explore the functional utility of our CD-promoter-based AAV vectors, we investigated the roles of molecular layer interneurons and Golgi cells in cerebellar functions, including motor control, emotion, and higher cognitive functions such as social behaviors. Although previous studies have implicated cerebellar molecular layer interneurons in social behavior ^17, 18^, their roles in other behaviors such as anxiety and motor functions are less understood. Moreover, the roles of Golgi cells in behaviors remain directly unexplored.

To examine these roles, we used AAV-RE16_CD68-M3Dq-mcherry to taget molecular layer interneurons and AAV-HGT017_CD9-DIO-M3Dq-mcherry to target Golgi cells, enabling chemogenetic activation via the hM3Dq receptor. For mice infected with AAV-RE16_CD68-M3Dq-mCherry, intraperitoneal injection of CNO significantly decreased the time spent in the center of the open-field arena compared with PBS injection in control mice (Figure. 5A), suggesting that activation of MLIs increased anxiety-like behavior. Additionally, CNO administration led to a significant increase in social interaction time in the three-chamber social test (Figure. 5B), indicating enhanced social behavior upon activation of MLIs. In the rotarod test, CNO injection improved motor performance compared with control mice (Figure. 5C), possibly due to enhanced motor learning. These findings provide new insights into the role of MLIs in modulating anxiety and motor functions, expanding our understanding beyond their previously known involvement in social behaviors

**Figure. 5:**
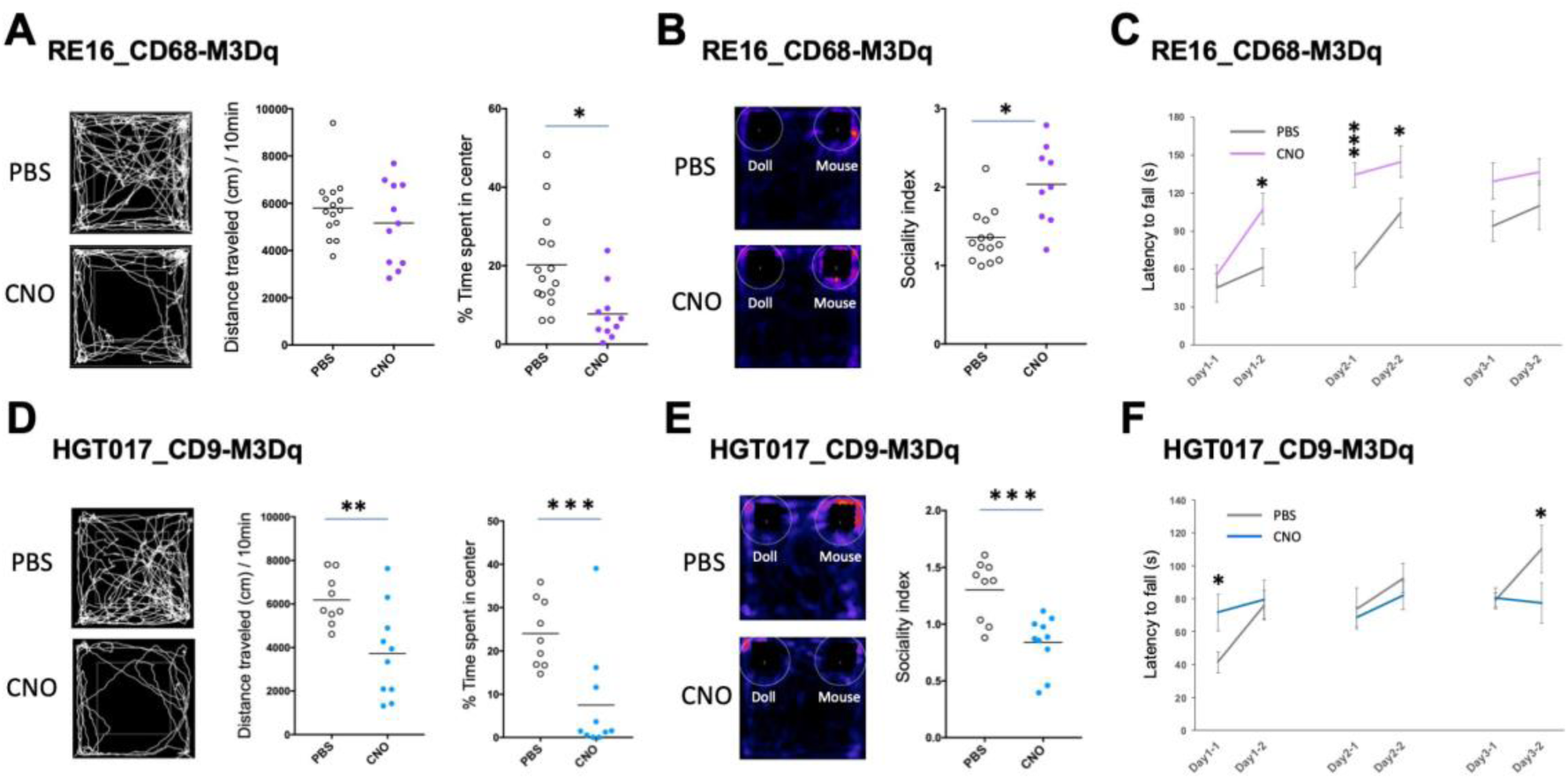
Modulation of behaviors by manipulation of specific cell types in the cerebellum. **A-C,** behavioral tests for mice infected by the AAV with mscRE16_hCD68-M3Dq-mcherry and the comparison between PBS injected mice (control) and CNO injected mice. **A**, open filed test. Representative tracks (left) and summary graphs showing the total distance during 10 min (middle) and % time spent in the center region of the box (right) for PBS control (white circles, n = 15) and CNO (purple circles, n = 11) mice. **B,** the sociality test. Representative heatmaps (left) and summary graphs showing the sociality index (right) for PBS control (white circles, n = 15) and CNO (purple circles, n = 11) mice. Sociality index = (Tmouse)/(Tdoll) where Tmouse and Tdoll are the staying time around the mouse cage and that around the doll cage, respectively. **C,** rota-rod test. The latency for falling off the rod is shown for PBS control (black lines, n = 15) and CNO (purple lines, n = 11) mice. **D-F,** panels similar to a-c for the AAV with HGT017_hCD9-M3Dq-mcherry. Data for PBS control (black lines, n = 9) and CNO (light blue lines, n = 10) mice are shown. *\*P* < 0.05, *\*\*P* < 0.01, *\*\*\*P* < 0.001 (Mann-Whitney *U* test). Data are mean ± SEM.

In mice infected with AAV-HGT017_CD9-DIO-M3Dq-mCherry, CNO treatment caused a decline in locomotor activity and heightened anxiety, as demonstrated by reduced distance traveled and less time spent in the center of the open-field arena compared with PBS injection in control mice (Figure. 5D). This suggests that activation of Golgi cells decreases locomotor activity and increases anxiety-like behavior. Moreover, CNO injection into these mice significantly decreased social interaction time (Figure. 5E), indicating a reduction in social behavior upon activation of Golgi cells. In the rotarod test, CNO administration initially enhanced motor performance at the first trial but decreased performance at the last trial compared with control mice (Figure. 5F), suggesting complex effects on motor coordination and learning. These results highlight the previously underexplored role of Golgi cells in modulating a range of behaviors, providing new insights into their functions.

### Rescue of sociality deficit in ASD model mice by a CD-promoter-based AAV

Since we found that CNO injection into mice with AAV-RE16_CD68-M3Dq-mCherry enhanced sociality (Figure. 5), we examined the therapeutic potential of CD-promoter-based AAV vectors in ameliorating social and motor deficits in an autism spectrum disorder (ASD) mouse model. Although previous studies have implicated cerebellar molecular layer interneurons in social behavior ^17, 18^, direct evidence demonstrating that targeted activation of these neurons can rescue social deficits in ASD models is lacking.

To generate ASD model mice and control mice, we injected AAVs carrying L7-iCre or L7-EGFP, respectively, into the cerebellum of tuberous sclerosis complex 1 (TSC1) floxed mice at postnatal day 3–4^19^. The TSC1 conditional knockout mice exhibit social and motor deficits resembling ASD phenotypes. At postnatal weeks 8–9, we administered AAV-RE16_CD68-M3Dq-mCherry into the cerebellum of these mice to enable selective activation of molecular layer interneurons. We compared behaviors among three groups of mice: (1) control mice (TSC1 floxed mice infected with AAV-L7-EGFP and AAV-RE16_CD68-M3Dq-mCherry, PBS injection before behavioral tests), (2) TSC1 conditional knockout mice (infected with AAV-L7-iCre and AAV-RE16_CD68-M3Dq-mCherry, PBS injection before behavioral tests), and (3) TSC1 conditional knockout mice with M3Dq activation (infected with AAV-L7-iCre and AAV-RE16_CD68-M3Dq-mCherry, CNO injection before behavioral tests). Our results confirmed that TSC1 conditional knockout mice exhibited a reduction in preference towards stranger mice compared to control mice^19^ (Figure. 6A-B). Importantly, M3Dq activation of molecular layer interneurons by CNO injection in TSC1 conditional knockout mice led to a significant improvement in social behavior compared to TSC1 conditional knockout mice treated with PBS (Figure. 6A-B). This suggests that activation of molecular layer interneurons can ameliorate social deficits in this ASD model, providing new evidence for their role in social behaviors. Additionally, Additionally, we attempted to rescue motor deficits observed in TSC1 conditional knockout mice. Using the accelerating rotarod test, we confirmed the motor function deficit in TSC1 conditional knockout mice (Figure. 6C). Activation of molecular layer interneurons via M3Dq activation by CNO significantly improved motor performance in TSC1 conditional knockout mice compared to those without M3Dq activation (Figure. 6C). These results suggest that activation of molecular layer interneurons ameliorates both social and motor deficits in TSC1 conditional knockout mice.

**Figure. 6:**
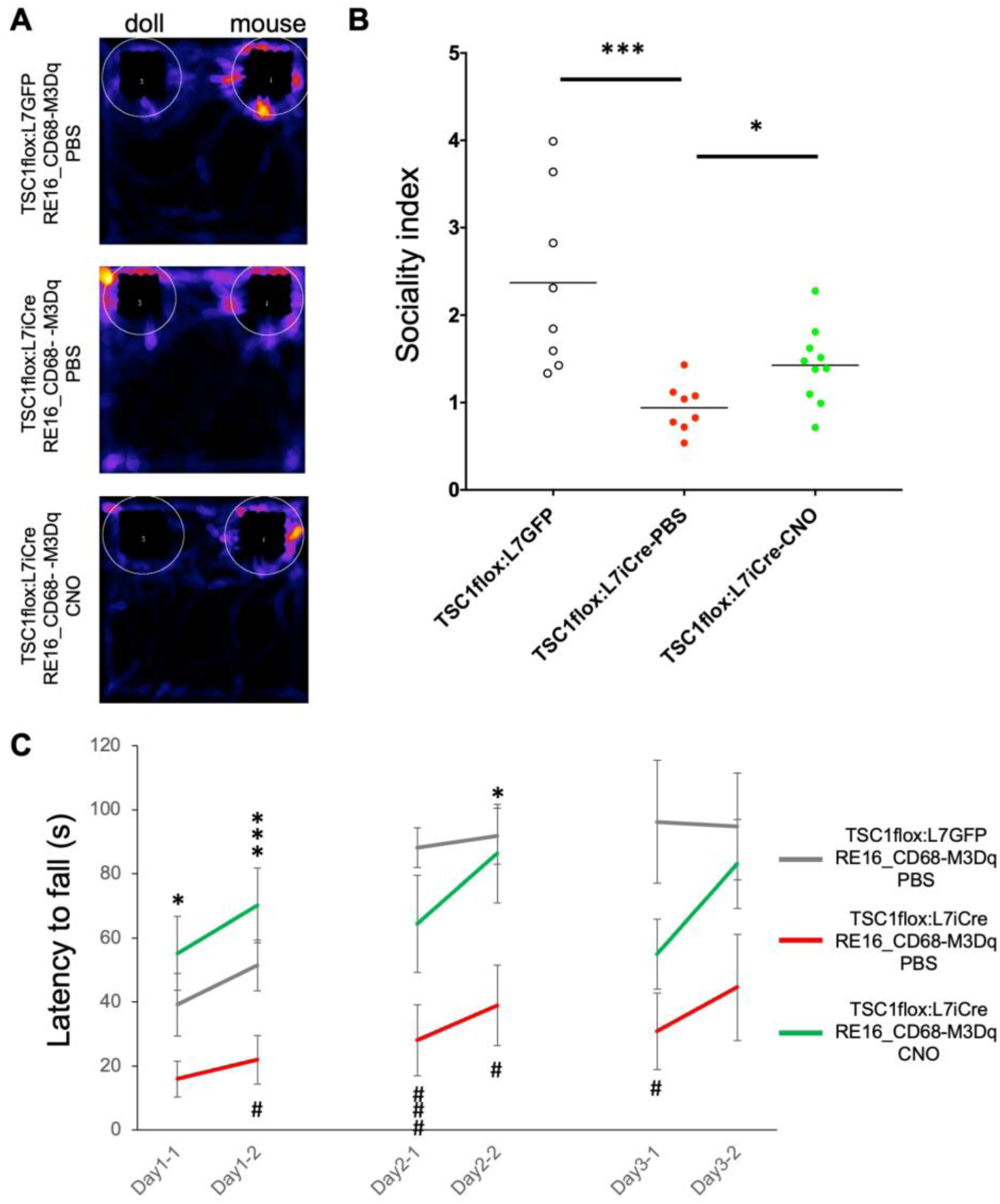
Rescue of sociality deficit in TSC1 mutant mice by CD-promoter-based AAV. **A,B,** the sociality test for control mice (TSC1 flox mice infected by AAV-L7-EGFP at P3-4 and AAV-RE16_CD68-M3Dq-mcherry at postnatal week 8-9, PBS application before behavior tests), TSC1 conditional knockout mice (those infected by AAV-L7-iCre at P3-4 and AAV-RE16_CD68-M3Dq-mcherry at postnatal week 8-9, PBS application before behavior tests), TSC1 conditional knockout mice with M3Dq activation (those infected by L7-iCre at P3-4 and RE16_CD68-M3Dq-mcherry at postnatal week 8-9, CNO application before behavior tests). **A**, representative heatmaps for three types of mice. **B**, summary graphs showing the sociality index for control (white circles, n = 8 mice), TSC1 conditional knockout (red circles, n = 8 mice), and TSC1 conditional knockout with M3Dq activation (green circles, n = 10 mice) mice. Sociality index = (Tmouse)/(Tdoll) where Tmouse and Tdoll are the staying time around the mouse cage and that around the doll cage, respectively. **C,** rota-rod test. The latency for falling off the rod is shown for control (black lines, n = 8 mice), TSC1 conditional knockout (red lines, n = 8 mice), and TSC1 conditional knockout with M3Dq activation (green lines, n = 10 mice) mice. # represents the statistically significant difference between L7GFP:RE16_CD68-M3Dq and L7iCre:RE16_CD68-M3Dq-PBS, * represents the significant difference between L7iCre:RE16_CD68-M3Dq-PBS vs L7iCre:RE16_CD68-M3Dq-CNO), *\* or ^#^ P* < 0.05, *\*\*\* or ^###^ P* < 0.001 (Mann-Whitney *U* test). Data are mean ± SEM.

## Discussion

In this study, we have successfully developed cell-type specific adeno-associated viruses (AAVs) with the Cluster of Differentiation (CD) promoters to target specific cell types in the mouse cerebellum and cortex. Our findings demonstrate the utility of CD-promoter-based AAVs for manipulating neuronal activity in a cell-type-specific manner, overcoming existing technical barriers by enabling targeted manipulation without the need for genetically engineered animals. By providing new insights into the roles of specific cell types such as MLIs and Golgi cells in various behaviors, our study advances the understanding of brain function and underscores the potential of our approach for neuroscience research and therapeutic strategies. One unique advantage of CD promoters is their potential to target neuronal subtypes that are challenging to label with existing promoters. Since CD genes are involved in cell differentiation and are expressed in various cell types, including neurons, they may offer alternative regulatory elements to achieve specific expression patterns.

Our CD-promoter-based AAVs demonstrated remarkable cell-type specificity. Specifically, the promoters of CD68 and CD9 exhibited high specificity in driving reporter gene expression in cerebellar interneurons. Notably, the AAV vector utilizing the CD9 promoter is the first to enable specific labeling of Golgi cells in the cerebellum. The level of specificity by CD68 promoter is comparable to that achieved by the *GAD65* promoter ^3^. Similarly, the CD1d and CD52 promoters in the cortex exhibited high specificity on par with the established promotes and enhancers like *GAD65* promoter ^3^, *Scn1a* enhancers ^6^, and the *CaMK2* promoter ^16^. Discovering new promoters with the same cell specificity allows for greater flexibility and optimization in gene therapy approaches. Different promoters may vary in their expression strength, regulation, and compatibility with various cell types or disease conditions. By expanding the repertoire of available promoters, researchers can select the most appropriate DNA element to achieve desired expression levels, temporal control, or to overcome limitations posed by the cellular environment in different disease models. Moreover, we further fine-tuned the cell type specificity by combining CD promoters with enhancer elements and miRNA targeting sequences. Such composite regulatory DNA sequences can significantly boost the precision of AAV-mediated gene manipulation in targeted cell types, enhancing the relevance and applicability of our approach.

In contrast to transgenic animals, our study advances the use of AAV vectors as a simpler and more cost-effective alternative. This innovation will accelerate the elucidation of the input-output relationships among distinct cell types, their respective roles in brain function and development, their interactive dynamics, and their implications for novel treatments for brain diseases. However, AAV vectors have a limitation. AAV vectors can accommodate only the limited size of genetic material. We utilized CD promoters ranging from 700bp to 2000bp in length. Notably, a 2000bp promoter is relatively long for AAV vector Packaging ^20^. Researchers must carefully consider the size and number of genes they wish to deliver when selecting an appropriate vector system that aligns with their research or therapeutic objectives. The limited size of genetic material can be a challenge when trying to deliver large genes or multiple genes in a single vector. Recent strategies have emerged to circumvent the size constraint, such as employing multiple vectors or fragmenting larger genes into smaller pieces for independent vector packaging and subsequent reassembly in the target cell ^21–23^.

Using our CD-promoter-based AAVs, we clarified the distinct and shared roles of molecular layer interneurons and Golgi cells in regulating cerebellar functions such as anxiety, sociality and motor control. The distinct roles may arise from cell types that each interneuron targets: Molecular layer interneurons can modulate cerebellar output directly by inhibiting Purkinje cells, whereas Golgi cells can suppress cerebellar input by inhibiting granule cells. Activation of molecular layer interneurons may enhance the reward value or salience of social stimuli and boost social motivation ^24, 25^. The increased anxiety by molecular layer interneuron activation may reflect a heightened alertness or arousal state that facilitates social and motor behaviors. Conversely, activation of Golgi cells might lead to increased anxiety and decreased locomotor activity, reflecting a state of reduced motivation and exploration that adversely impacts social and motor behaviors.

We demonstrates that activation of molecular layer interneurons in the cerebellum ameliorates the sociality deficit in ASD model mice. This finding aligns with previous studies that implicate cerebellar dysfunction in the pathophysiology of ASD ^19, 26, 27^. It is possible that activation of molecular layer interneuron restores the balance of excitation and inhibition in the cerebellum, thereby enhancing the communication with other brain regions crucial for social behaviors ^24, 28–30^. It is worth noting that the specificity of the CD-promoter-based AAV may vary between different animal models and human patients. Further optimization and validation steps are necessary before any clinical application can be considered.

We provide novel insights by demonstrating that targeted activation of cerebellar molecular layer interneurons using our CD-promoter-based AAV vector significantly ameliorates social deficits in ASD model mice. While previous studies have implicated these interneurons in social behaviors ^17, 18^, our work offers direct evidence of their therapeutic potential. Although the mechanisms by which activation of molecular layer interneurons rescues ASD-like behaviors remain unknown, the activation may restore the excitation-inhibition balance within cerebellar circuits, thereby enhancing communication with other brain regions critical for social functioning (Carta et al., 2019; D’Mello and Stoodley, 2015; Kelly et al., 2020; Stoodley and Tsai, 2021). This finding advances our understanding of the cerebellum’s role in ASD by identifying a precise cellular mechanism that can be targeted for intervention. Moreover, our use of the CD-promoter-based AAV vector introduces a novel gene therapy approach with enhanced cell-type specificity compared to existing vectors. This specificity is crucial for minimizing off-target effects and maximizing therapeutic efficacy in gene therapy applications. However, we acknowledge that the specificity of the CD-promoter-based AAV may vary between different animal models and human patients, necessitating further optimization and validation before clinical application can be considered.

Additionally, our work expands the utility of CD promoters by demonstrating their scalability in targeting specific cell types in the cerebellum and cortex. CD genes, primarily known as cell-surface molecules on leukocytes and other immune cells, possess broad applicability given the diversity of CD antigens (over 370 CD antigens identified in humans). Since CD genes are also expressed in other organs such as blood cells and immune cells, AAVs utilizing CD promoters are not only useful in neuroscience but also have implications in many fields including immunology and hematology.

In conclusion, we have engineered a series of AAVs with CD promoters that can precisely target specific populations of cells. These CD promoters-based AAV vectors can be further customized through the incorporation of various enhancers or miRNA targeting sequences. Our work paves the way to access and manipulate specific cell types for both basic and translational research.

## Materials and Methods

### Animals

This study was conducted under the recommendations of the Regulations for Animal Experiments and Related Activities at Tokyo Medical and Dental University. All animal experiment was approved by the Committees on Gene Recombination Experiments and Animal Experiments of Tokyo Medical and Dental University. All animals were group housed and maintained on a 12hr:12hr light dark cycle. All efforts were made to reduce the number of animals used and to minimize the suffering and pain of animals. WT mice (ICR and C57BL6N, 8-16 weeks, Sankyo labo service) and TSC1 flox mice (postnatal day 3 to postnatal week 10, Jackson lab, Strain# 005680) were used in this study.

### Plasmids

The CD gene promoter sequences were amplified by PCR from mouse and human genomic DNA using primer pairs (Table. 1) and inserted into sites of hSyn promoter in AAV-U6-sgRNA-hSyn-mCherry (a gift from Alex Hewitt, Addgene plasmid # 87916) and in AAV-syn-EGFP (a gift from Bryan Roth, Addgene plasmid # 50465). hSyn promoter was removed by ApaI and BaMH1 or XbaI. The promoters, enhancers, and reporters were cloned using the Gibson Cloning Assembly Kit (New England BioLabs) following standard procedures. Specifically, for pAAV-L7-6-iCre, we amplified the iCre coding sequence from the pCAG-iCre (a gift from Wilson Wong, Addgene plasmid # 89573) and cloned it into pAAV/L7-6-GFP-WPRE (a gift from Hirokazu Hirai, Addgene plasmid # 126462); for AAV-S5E6 and AAV-HGT017, we amplified these sequences from the pAAV-S5E6-dTom-nlsdTom (a gift from Jordane Dimidschstein, Addgene plasmid # 135641) and the CN1253-scAAV-eHGT_017h-minBG-SYFP2-WPRE3-BGHpA (a gift from The Allen Institute for Brain Science & Boaz Levi, Addgene plasmid # 163497), and cloned them into 5’ region of CD9 promoter; for AAV-mscRE16, we amplified the sequences from the AiP1002-pAAV-mscRE16-minBGpromoter-EGFP-WPRE-hGHpA (a gift from The Allen Institute for Brain Science & Bosiljka Tasic, Addgene plasmid # 163486), and cloned it into 5’ region of CD68 promoter. For AAV-M3Dq-mcherry, we amplified the hM3D(Gq)-mCherry sequence from the pAAV-hSyn-DIO-hM3D(Gq)-mCherry (a gift from Bryan Roth, Addgene plasmid # 44361) and cloned it into the pAAV-mscRE16-CD68 and pAAV-HGT017-CD9. For pAAV-HGT017-CD9-mcherry-miR_targeting_sequences (Table. 2), four copies of single miR_targeting sequence were amplified and inserted between the 3’ end of the mcherry insert and the 5’ end of hGH polyA regulatory sequence insert in pAAV-HGT017-CD9-mcherry. For AAV with jGCaMP8s, the mcherry sequence was removed from pAAV-HGT017-CD9-mcherry-miR133a_targeting and the cDNA of jGCaMP8s was inserted into the removed mcherry region of pAAV-HGT017-CD9-mcherry-miR133a _targeting.

### AAV packaging and injection

The AAVs were produced using standard production methods. HEK293 cells were transfected with by Polyethylenimine. AAV9, PHPeB^13^, and CAP-B10^14^ were used. Virus was collected after 120 h from both cell lysates and media. Purification Kit (Takara) was used for viral particle purification. All batches produced were in the range 10^10^ to 10^11^ viral genomes per milliliter.

The procedure for viral vector injection has been modified from the previous lentivirus protocol^11^. Animals were anesthetized with a mixture of midazolam (4 mg/kg body weight (BW)), butorphanol (5 mg/kg BW), and medetomidine (0.3 mg/kg BW). The depth of anesthesia was constantly monitored by observing pinch-induced forelimb withdrawal reflex. 1-1.5 µL of viral solution were injected into the cerebellar vermis and the S1 cortex of adult C57BL/6N mice at 100 nl / min. 1.5 µL of pAAV-L7-6-iCre and pAAV-L7-6-GFP was injected into the cerebellum of TSC1 flox mice at postnatal day 3-4, and then 1.5 µL of pAAV-mscRE16-CD68-M3Dq-mcherry was injected into the cerebellum at postnatal week 8-9. For retro-orbital injection, 100 μL of the viral solution was intravenously injected into the retro-orbital sinus of mice using a 0.5 ml syringe with a 30-gauge needle (326668; Becton Dickinson, Japan).

### Immunohistochemistry

Mice were perfused with 4% paraformaldehyde in 0.1 M phosphate buffer, and processed for parasagittal microslicer sections (100 μm in thickness). After permeabilization and blockade of nonspecific binding, the following antibodies were applied for 2 days at 4 °C: an antibody against Car8 (Car8-GP-Af500 or Car8-Go-Af780, diluted 1:300, Nittobo Medical) to visualize PCs; an antibody against S100b (S100b-GP-Af630, 1:300, Nittobo Medical); an antibody against parvalbumin (PV-Go-Af460, 1:300, Nittobo Medical), an antibody against RFP (390004, 1:1000, Synaptic Systems or PM005, 1:500, MBL), an antibody against neurogranin (AB5620, 1:1000, Merck Millipore), an antibody against CaMK2 (ab52476, 1:500, abcam), an antibody against NeuN (MAB377, 1:1000, Merck Millipore), and/or an antibody against GFP (#06083-05, 1:1000, Nacalai Tesque). After incubation with secondary antibodies (an anti-rat Alexa Fluor 488, an anti-guinea pig Cy3, an anti-goat Alexa Fluor 488, an anti-goat Alexa Fluor 647, an anti-mouse Cy5, an anti-rabbit Cy3, an anti-rabbit Alexa Fluor 488), the immunolabeled sections were washed and then examined under a fluorescent microscope (BZ-X700 or BZ-X800, Keyence).

### In vivo calcium imaging

Animals were put on a warm blanket and anesthetized with a mixture of midazolam (4 mg/kg body weight (BW)), butorphanol (5 mg/kg BW), and medetomidine (0.3 mg/kg BW). The depth of anesthesia was constantly monitored by observing pinch-induced forelimb withdrawal reflex. The skin and muscles over the skull were removed and a metal plate was fixed with dental acrylic cement over the lobule 5-7 of the cerebellar vermis. A craniotomy of 3 mm diameter was made and the dura mater was carefully removed. A sterile circular 3mm diameter glass coverslip was put directly on the dura mater and was secured in place with surgical adhesive (Alon-Alpha A, Sankyo). A custom-made metal headplate with a 5 mm circular imaging well was fixed to the skull over the cranial window with dental cement (Super-Bond, Sun-medical, Japan). In vivo calcium imaging was performed using a two-photon microscope (Nikon, AX R MP) equipped with a ×16 objective lens (Nikon, CFI75 LWD 16X W) and a ultrafast laser (Axon 920-2 TPC). During the recordings, the animals were lied on a warm blanket to keep the body temperature at 37°C. Laser power was maintained <20 mW at the sample. Images were obtained in a 2D plane (512 × 512 pixels, 890 µm x 890 µm) using Nikon NIS-Elements software (frame rate, 1 Hz).

To extract the fluorescence changes corresponding to the calcium transients, the raw movie was first converted to *Δ*F / Fb wave ^31^. *Δ*F / Fb was expressed as (F - Fb) / (Fb), where Fb is the baseline fluorescence in the absence of calcium transients. Fb was defined as follows. First, we calculated a threshold as mean + 1 SD of the F wave of each pixel, and then the F wave below the threshold was averaged to obtain Fb. The *Δ*F / F wave of each pixel was then high-pass filtered at 0.1 Hz to eliminate slow drifts presumably due to photo-bleaching effects or glial signals.

### Behaviors

Male mice from 2 to 3 months of age 7-15 days after AAV injection were used for behavioral tests. Before the experiments, the mice were habituated in the testing areas for at least 30 min. Mice behavior was recorded through the video camera set in the experimental room. DREADD agonist, CNO (Cat. No. 4936, Tocris, UK) was used to selectively activate the excitatory DREADD, hM3Dq. 1 mg/kg CNO was delivered intraperitoneally to each mouse 40 min before a behavioral test. The open field test was performed in a 50 cm x 50 cm x 40 cm (W x D x H) open field box for 10 min. Total distance mice traveled and the time in the center and peripheral regions were automatically analyzed by the ImageJ software (MouBeAT) ^32^.

The sociality test was performed in the open field apparatus (50 x 50 x 40 cm, W x D x H). The test consisted of a 10-min habituation session and a 10-min test session. In the habituation session, mice were allowed to explore the open field freely. In the test session, two quadrangular cages with slits (8 cm x 8 cm, 18 cm high) were placed in two adjacent corners. One cage contained a novel mouse (8-week-old male C57BL6N) and the other was a mouse doll, and a test mouse was first placed in the outer regions of open field arena at the longest distance from these cages. The times spent around the two cages areas (12 cm in diameter) were analyzed automatically with the ImageJ software (MouBeAT). Social preference was assessed by the following equation: Sociality index = (T_mouse_)/(T_doll_) where T_mouse_ and T_doll_ are the staying time around the mouse cage and that around the doll cage, respectively.

To assess motor coordination and motor learning, mice were subjected to a rotarod test. Mice were placed on a resting state rotarod (model LE8205, Panlab) for three consecutive days with two trials per day spaced 10 min apart (30 min and 40 min after CNO injection). For each trial, the rotarod was accelerated linearly from 4 rpm to 40 rpm over 300 s. Latency to fall from the start of rotation was measured.

Grooming test was performed in a cylindrical cage. Mice were placed in a new empty cage without bedding. The total recording time was 10 min. The spontaneous grooming behavior was recorded through the video camera set up in the experimental room. The footage was then analyzed.

### Statistical analysis

All data are presented as mean ± SEM. Statistical significance was assessed by Mann-Whitney U test for the comparison of two independent samples. To compare the two different categorical independent samples on one dependent variable, Two-way ANOVA with post-hoc test (Bonferroni correction) was used as indicated in the text. Statistical analysis was conducted with GraphPad Prism program. Differences between groups were judged to be significant when p values were smaller than 0.05. *, **, ***, **** and ns represents p < 0.05, p < 0.01, p < 0.001, p < 0.0001 and not significant, respectively.

## Data and Materials Availability

Data and Plasmids generated in this study are available upon reasonable requests. Further information and requests for resources and reagents should be directed to and will be fulfilled by the Lead Contact, Naofumi Uesaka.

## Acknowledgements

We thank Daisuke Tanaka for helpful discussions. This work was supported by Grants-in-Aid for Scientific Research (22H05092, 22H05093 to N.U.) from JSPS, Japan, by the Asahi Glass Foundation, by Takeda Science Foundation, and by Tokumori Yasumoto Memorial Trust for Researches on Tuberous Sclerosis Complex and Related Rare Neurological Disease, Japan.

## Author contributions

N.U. designed the study. N.U, M.S, R.M. and A.T. performed the experiments and/or data analysis. N.U. wrote the manuscript, which was reviewed by all authors.

