## Supplementary Figs 1-4 for "Viral vectors with cluster of differentiation gene promoters to target specific cell types in the brain"

Supplementary information includes Figure S1-S4.

**Fig. S1: CD promoters drive gene expression in specific type of cells in the mouse cerebellum.**

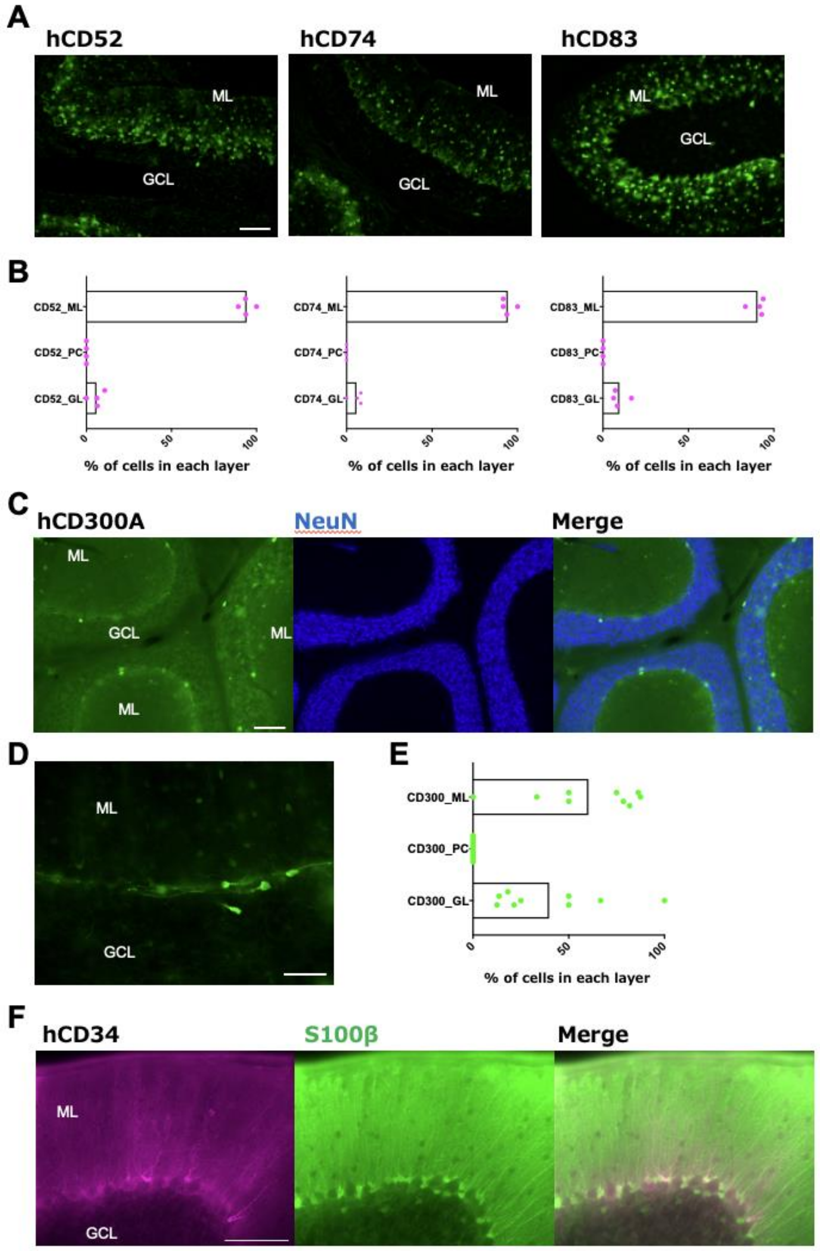


**A**, EGFP fluorescent images of the cerebellum infected by AAVs with one of hCD52, hCD74, and hCD83 promoters. **B**, graphs showing the percentage of $EGFP^+$ cells in each cerebellar layer for hCD52, hCD74, and hCD83 promoters. n = 2 mice for each. **C**, images showing EGFP fluorescence and NeuN staining of the cerebellum infected by the AAV with hCD300A promoter. **D**, a higher magnification image of EGFP labeled cells for hCD300A promoter. **E**, a graph showing the percentage of $EGFP^+$ cells in each cerebellar layer for hCD300A promoter. n = 2 mice. **F**, images showing mcherry fluorescence and S100β staining of the cerebellum infected by the AAV with hCD34 promoter. Scale bar, 100μm for a, c, 50μm for d, f. ML, molecular layer. PC, Purkinje cell. GL or GCL, granule cell layer.

**Fig. S2: Layer distribution of cells labeled by AAVs with mcherry under hSyanpsin promoter in the mouse cerebellum.**

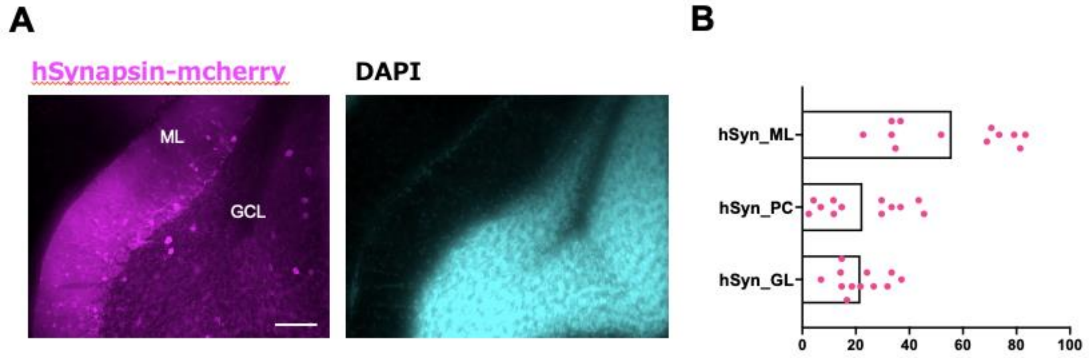


**A**, fluorescent images of mcherry (magenta) and DAPI (cyan) in the cerebellar slice infected by the AAV with the hsynapsin. Scale bar, 100 µm. **B**, the graph showing the percentage of mcherry$^{+}$ cells in each cerebellar layer. n = 3 mice. ML, molecular layer. PC, Purkinje cell. GL or GCL, granule cell layer.

**Fig. S3: mscRE16 Enhancer and CD68 promoter with the specificity to molecular layer interneurons in the mouse cerebellum.**

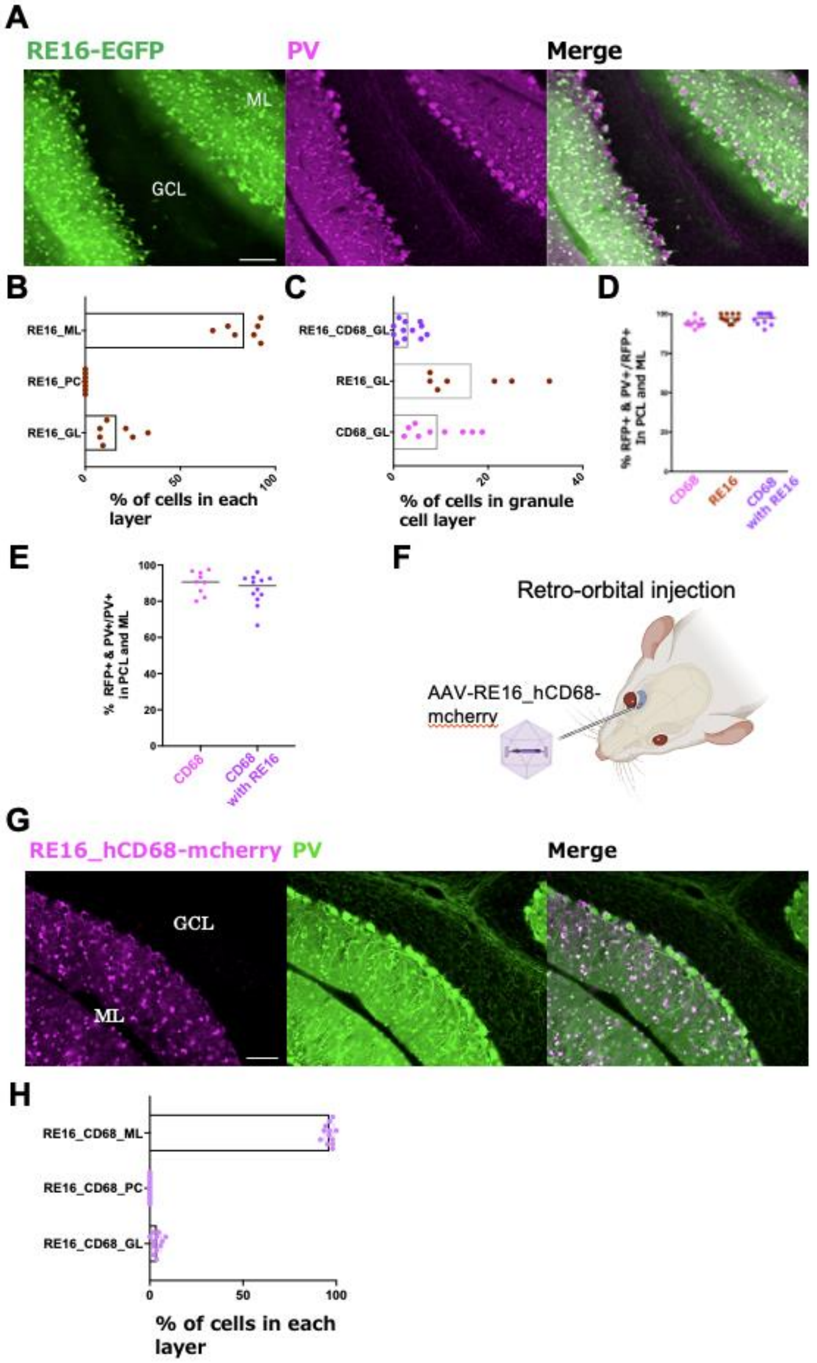


**A**, fluorescent images of EGFP (green) and PV (magenta) in the cerebellar slice infected by the AAV with the mscRE16 enhancer. **B**, the graph showing the percentage of $EGFP^+$ cells in each cerebellar layer for the mscRE16 enhancer. n = 2 mice. **C**, the graph showing the comparison of the percentage of $mcherry^+$ (hCD68) or $EGFP^+$ (mscRE16) cells in the granule cell layer among the mscRE16 enhancer, the hCD68 promoter, and the hCD68 promoter with the mscRE16 enhancer. **D**, the graph showing specificity of gene expression in $PV^+$ molecular layer interneurons for the mscRE16 enhancer, the hCD68 promoter, and the hCD68 promoter with the mscRE16 enhancer.

The percentages were determined by the ratio of double-labeled cells ($PV^+$ and $mcherry^+$ or $EGFP^+$) to total $mcherry^+$ or $EGFP^+$ cells. **E**, the graph showing infection efficacy of gene expression in $PV^+$ molecular layer interneurons for the hCD68 promoter, and the hCD68 promoter with the mscRE16 enhancer. The percentages were determined by the ratio of double-labeled cells ($PV^+$ and $mcherry^+$) to total $PV^+$ cells. **F**, Schematic of the retro-orbital injection procedure used to deliver AAV-RE16-hCD68-mcherry to the mouse brain. **G**, images (mcherry (magenta) and PV (green)) for mice with the retro-orbital injection of the AAV-RE16-hCD68-mcherry. **H**, the graph showing % of $mcherry^+$ cells in each cerebellar layer for the AAV-RE16-hCD68-mcherry. n = 2 mice. Scale bar, 100µm for A, G. ML, molecular layer. PC, Purkinje cell. GL or GCL, granule cell layer.

**Fig. S4: miRNAs and CD9 promoter with the specificity to granule cell layer interneurons in the mouse cerebellum.**

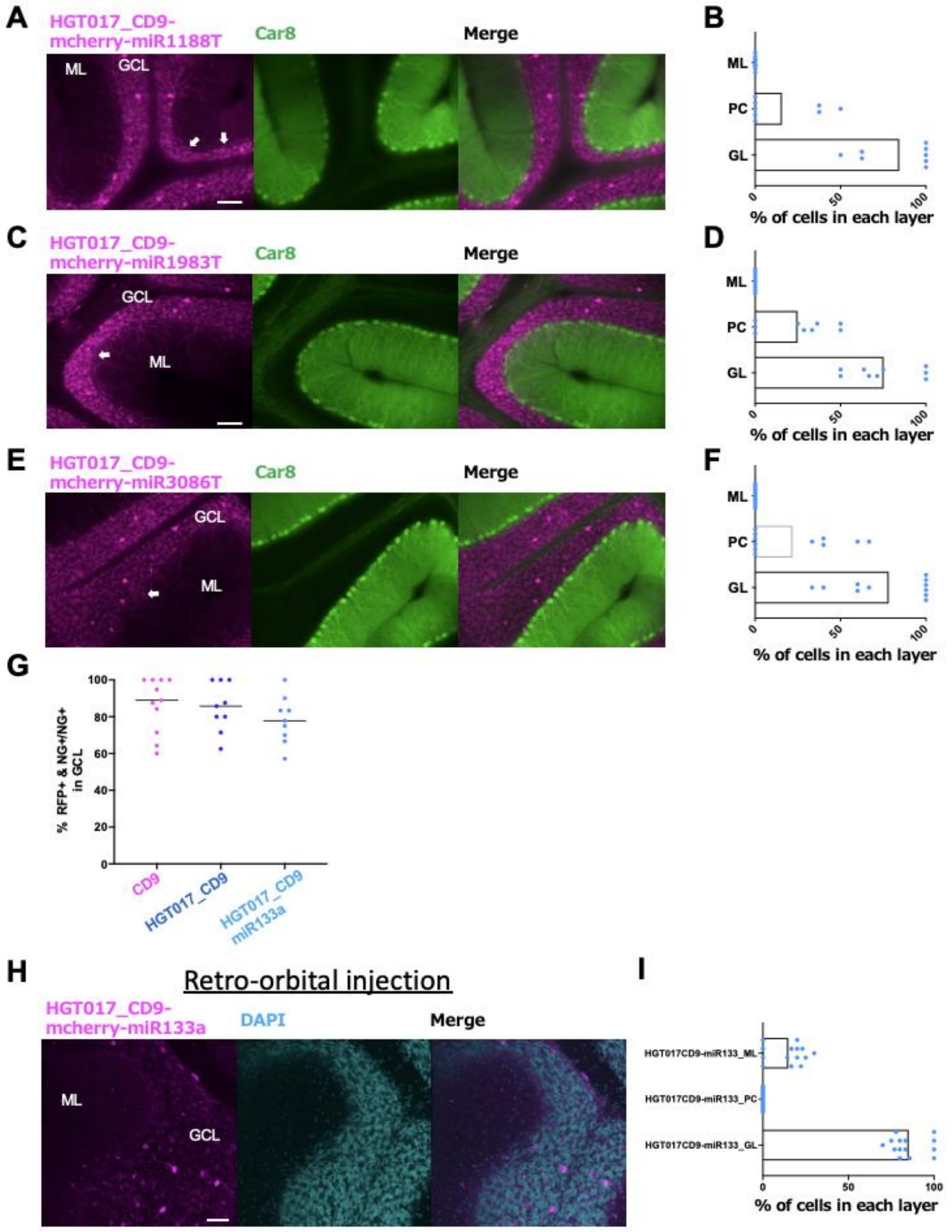


**A, C, E,** fluorescent images of mcherry (magenta) and Car8 (green) in the cerebellar slices infected by AAVs with mcherry under the hCD9 promoter with HGT017 enhancer and miRNA targeting sequences (A for miRNA1188, C for miRNA1983, E for miRNA3086). Arrows show mcherry-labeled Purkinje cells. **B, D, F**, graphs showing the percentage of mcherrys$^{+}$ cells in each cerebellar layer for AAVs with each miRNA targeting sequence and mcherry under the hCD9 promoter with HGT017 enhancer. n = 2 mice for each. **G**, the graph showing infection efficacy of gene expression in neurogranulin$^{+}$ granule layer interneurons for the hCD9 promoter, the hCD9 promoter with HGT017 enhancer, and the hCD9 promoter with the HGT017 enhancer and miRNA133a. The percentages were determined by the ratio of double-labeled cells (neurogranin$^{+}$ and mcherry$^{+}$) to total neurogranin$^{+}$ cells. **H**, images (mcherry (magenta) and DAPI (cyan)) for mice with the retro-orbital injection of the AAV-HGT017_CD9-mcherry-miR133a. **I**, the graph showing % of mcherry$^{+}$ cells in each cerebellar layer for the AAV-HGT017_CD9-mcherry-miR133a. n = 2 mice. Scale bar, 100 µm for A, C, E and 50 µm for H. ML, molecular layer. PC, Purkinje cell. GL or GCL, granule cell layer.
